# Oncolytic Maraba virus MG1 mediates direct and natural killer cell-dependent lysis of Ewing sarcoma

**DOI:** 10.1101/2025.05.16.654440

**Authors:** Tyler Barr, Victoria A. Jennings, Elizabeth A. Roundhill, Richard T. Baugh, Maisa Yamrali, Heather Owston, Dennis McGonagle, Peter V. Giannoudis, Natasha J. Caplen, Javed Khan, John C. Bell, Susan A. Burchill, Fiona Errington-Mais, Graham P. Cook

## Abstract

Ewing sarcoma (EWS) is a rare cancer of the bone and soft tissue, most prevalent in children and young adults. Treatment of EWS has progressed relatively little in over 30 years. Survival rates for patients, particularly those with metastatic and/or relapsed disease, remain poor, highlighting the urgent need for innovative treatment options. We have explored the therapeutic potential of the oncolytic Maraba virus strain MG1. We show that MG1 undergoes productive replication and exerts direct oncolysis of established EWS cell lines, doxorubicin-resistant EWS cell lines and patient-derived Ewing sarcoma (PDES) cell cultures more recently established from tumours. In contrast, primary mesenchymal stem cells (the likely cell of origin of EWS) were resistant to MG1, with IFN-I being a major determinant of tumour cell selectivity. MG1 treated PBMC produced IFN-I and killed EWS cells *in vitro,* in a natural killer (NK) cell-dependent manner. The ability of MG1 to kill EWS cells directly and to stimulate NK cell cytotoxicity against this tumour suggests that Maraba virus MG1 may provide therapeutic benefit for EWS patients where the efficacy of conventional treatments is currently limited.

## Introduction

Ewing sarcoma (EWS) was first described by James Ewing in 1921 (Ewing, 1921). This primary bone cancer predominantly affects children and young adults and is characterised by non-random chromosomal translocations between *FET* and *ETS* gene family members, which result in the expression of a fusion oncoprotein (Delattre et al., 1992; Sorensen et al., 1994). Approximately 85% of EWS cases involve a t(11;22)(q24;q12) translocation, generating an *EWSR1::FLI1* fusion oncogene, while ∼10% have t(21;22)(q22;q12) resulting in an *EWSR1::ERG* fusion oncogene, a number of rarer translocations have also been documented (Dupuy et al., 2023; Grünewald et al., 2018; Le Deley et al., 2010; Sorensen et al., 1994).

Treatment for EWS has changed very little in the last three decades, with combinations of chemotherapies (vincristine, ifosfamide, doxorubicin, etoposide and cyclophosphamide), radiotherapy and surgery being used (Brennan et al., 2020; Zöllner et al., 2021). Currently, approximately 65-75% of patients with localised disease at diagnosis survive for five years or more (Gaspar et al., 2015). However, patients with metastatic or relapsed disease have a much poorer prognosis, with a 5 year survival rate of less than 30% (Gaspar et al., 2015). This highlights a need for new treatment approaches, especially for patients with disseminated or relapsed/refractory disease.

The EWSR1::FLI1/ERG fusion oncoproteins function as aberrant transcription factors and drive many of the key features of EWS by acting as potent modulators of gene expression (Bailly et al., 1994; Guillon et al., 2009; Hancock & Lessnick, 2008; Prieur et al., 2004). Furthermore, these fusion oncoproteins provide a highly selective drug target and a small molecule (termed TK-216 or derivative YK-4-279) has been developed which disrupts the interactions between EWSR1::FLI1 and its binding partner, RNA helicase A. This drug demonstrated strong activity in preclinical EWS models (Barber-Rotenberg et al., 2012). However, a recent phase I/II clinical trial reported limited efficacy in a cohort of relapsed/refractory EWS patients, highlighting a continued need for alternative treatment approaches (Meyers et al., 2024).

Immune checkpoint inhibitors have revolutionised cancer immunotherapy and agents targeting programmed cell death protein-1 (PD-1), programmed death-ligand 1 (PD-L1) and cytotoxic T-lymphocyte associated protein-4 (CTLA-4) are approved for use in many solid tumour types (Sun et al., 2023). Malignancies such as melanoma and non-small cell lung cancer, with high mutational burden and neoantigen load, generally respond well to immune checkpoint-based therapies, however, these are much less effective in EWS (Spurny et al., 2018; Sun et al., 2023; Tawbi et al., 2017; Thanindratarn et al., 2019). This is likely due to the low mutational burden in EWS, which generates few neoantigens and hence relatively weak T cell responses compared to tumours with a high neoantigen load (Crompton et al., 2014).

Despite disappointing results for immune checkpoint-based therapies, there is mounting evidence of anti-tumour immune responses in EWS, supporting the idea that immunotherapeutic approaches may hold promise for patients. Cytotoxic lymphocytes (CD8+ T cells and natural killer [NK] cells) are present in the EWS tumour microenvironment (TME) and their presence is associated with better overall survival (Berghuis et al., 2011; Cillo et al., 2022; Kuo et al., 2025; D. Stahl et al., 2019; Visser et al., 2023). Moreover, rates of lymphocyte recovery post-chemotherapy treatment are associated with improved patient prognosis (De Angulo et al., 2007). This indicates that EWS patients may indeed benefit from the application of immunotherapies, but different approaches may be required.

Oncolytic viruses (OV) are a cancer immunotherapy which target tumours via two broad mechanisms, direct oncolysis associated with the release of progeny virions and the activation of anti-tumour immunity (Jhawar et al., 2017). Moreover, OV can reprogramme the TME by induction of pro-inflammatory cytokines and chemokines, reducing immunosuppression and generating an immunologically “hot” TME (Armstrong et al., 2024). The preference of OV for cancer cells is not completely understood but is associated with the frequent inability of tumour cells to mount effective antiviral responses, thus making them permissive for viral replication (Aref et al., 2020; Brun et al., 2010; Müller et al., 2020). By contrast, OV-mediated production of type I interferon (IFN-I) by healthy cells stimulates innate immunity, in particular the activation of NK cells, which can detect and destroy both virally infected and malignant cells, placing them in a central position to mediate responses to OV infection of tumour cells (Parrish et al., 2015; Wantoch et al., 2022).

The efficacy of OV against EWS has been previously reported for a range of molecularly distinct OV, including protoparvovirus H-1, herpes simplex virus (rRp450), adenovirus (XVir-N-31), reovirus, vaccina virus, measles virus and vesicular stomatitis virus (Abdelbary et al., 2014; Eshun et al., 2010; Klose et al., 2019; Lacroix et al., 2018; Le Boeuf et al., 2017). However, EWS cells are often included amongst a panel of other sarcoma types, and detailed studies focussing on OV and EWS have not been undertaken. Importantly, EWS cell lines can be recognised and killed by NK cells, but efficient killing requires NK cell activation by exogenous agents such as IL-2 or IL-15 (Holmes et al., 2011; Pahl et al., 2012; Verhoeven et al., 2008). We have previously demonstrated that OV activate human NK cells both *in vitro* and *in vivo*, suggesting that OV might provide a useful stimulus to recruit NK cell activity against EWS (El-Sherbiny et al., 2015; Parrish et al., 2015; Wantoch et al., 2022).

Here, we have investigated the rhabdovirus, Maraba virus strain MG1, as a potential OV therapy for EWS in a range of human model systems. Maraba virus is a (+) single stranded RNA (ssRNA) virus originally isolated from sandflies but, like other rhabdoviruses, has the capacity to replicate in mammalian cells (Brun et al., 2010). Mutations in the virus M protein (L123W) and G protein (Q242R) were introduced to improve tumour cell selectivity, which is partially IFN-I dependent; this variant has the designation MG1 (Brun et al., 2010). MG1 has direct oncolytic activity against many cancer cell lines derived from several tumour types (Bourgeois-Daigneault et al., 2016; Brun et al., 2010; Hassanzadeh et al., 2019). Here, we describe the effects of MG1 on direct and immune-mediated killing of EWS using established cell lines (cultured as both monolayers and spheroids), doxorubicin-resistant EWS cell lines and patient-derived Ewing sarcoma (PDES) cell cultures, recently isolated from patient tumours. Our results demonstrate that MG1 exhibits efficient direct oncolysis and activates NK cell-dependent immune-mediated killing, suggesting that MG1 might provide a valuable therapeutic option for EWS.

## Materials and Methods

Details of cell lines, primary cultures including growth medium are provided in Table S1. Details of other reagents including antibodies, fluorescent stains and buffers used in this study are detailed in Table S2.

### Cell culture

All cell cultures were routinely tested for mycoplasma using Lookout® One-Step Mycoplasma PCR Detection Kit (Sigma-Aldrich) and were free from contamination. Unless otherwise stated, all growth medium was supplemented with 10% foetal bovine serum (FBS; Sigma), which was heat inactivated at 56°C for 30 minutes before use.

#### Cell lines

EWS cell lines SK-N-MC, TC-32, TTC-466, SK-ES-1 were studied. All cell lines were short tandem repeat (STR) profiled to validate their authenticity using ATCC STR profiling service.

#### Mesenchymal stem cells

Bone marrow derived mesenchymal stem cells (MSCs) were isolated from human healthy donor bone marrow aspirates. Ethical approval was obtained from NREC Yorkshire and Humberside National Research Ethics Committee (18/YH/0166). Isolation and analysis of MSC was as previously described (Wilson et al., 2024).

#### Peripheral blood mononuclear cells

Peripheral blood mononuclear cells (PBMC) were obtained from healthy donor leukocyte apheresis cones supplied by the National Health Service Blood and Transplant unit (NHSBT). PBMCs were isolated using Lymphoprep^TM^ (STEMCELL Technologies) and density gradient centrifugation.

#### Patient-derived EWS (PDES) cell cultures

PDES cell cultures were isolated from treatment naïve EWS patient biopsy tissue, as previously described (Roundhill et al., 2019), validation of EWS phenotype including fusion status and CD99 expression have been previously described (Roundhill et al., 2021). Informed consent and ethical approval for the collection of the tumours and generation of PDES cell cultures was obtained through GenoEWING (IRAS 167880, EDGE 79301).

#### Doxorubicin-resistant EWS cell lines

Doxorubicin-resistant cell lines were generated by maintaining SK-N-MC and TC-32 cell lines in half maximal effective concentration (EC50) of doxorubicin as determined by trypan blue exclusion for up to 6 months, and were then re-challenged with the 10-fold EC50 of doxorubicin until the EC50 population was stable, as previously described (Roundhill et al., 2019).

#### Spheroid cultures

Three-dimensional (3D) spheroid cultures were generated by seeding 2×10^3^ EWS cells into a 96 well U bottom ultra-low adhesion plate (Corning) for 7 days. Half media changes were carried out every 2-3 days.

### Oncolytic Maraba virus MG1

MG1-GFP and MG1-firefly luciferase (MG1-FLUC) were amplified on Vero cells. Standard plaque assay on Vero cell line was used to assess the viral titre of MG1 stocks and MG1-treated cell lysates. The virus backbone has previously been described (Brun et al., 2010).

### Assessment of cell viability and growth inhibition

Cell viability was evaluated using LIVE/DEAD® Fixable Yellow Dead Cell Stain Kit (Thermofisher). Growth inhibition of monolayer cultures was measured using Methylthialazole Tetrazolium (MTT; Sigma) assay and CellTiter-Glo® reagent (Promega) for spheroid cultures. All assays were conducted as per manufacturers’ instructions.

### NK cell depletion and flow cytometry-based functional assays

NK cells were depleted from whole PBMC using CD56 magnetic beads (Miltenyi Biotec). Briefly, PBMC were labelled with CD56 beads in MACS buffer for 15 minutes. Cells were then passed through LS columns (Miltenyi Biotec), mounted on a magnet, and washed 3 times with MACS buffer. Flow through was collected, and CD56+ NK cell depletion was validated using anti-CD3 and anti-CD56 antibodies and flow cytometry.

For immune-killing assays, whole or NK cell-depleted PBMC were treated with MG1 for 48 hours. PBMC were then co-cultured with CellTracker™ Green CMFDA Dye (Thermofisher)-stained EWS targets at a ratio of 25:1 for 5 hours. PBMC-EWS co cultures were then stained with LIVE/DEAD™ Fixable Yellow Dead Cell Stain Kit (Thermofisher).

For NK cell degranulation assays, PBMC were co-cultured with targets at a ratio of 10:1 for 1 hour, before the addition of anti-CD3, anti-CD56 and anti-CD107a antibodies and Brefeldin A at a final concentration of 3 µg/mL (Table S2). Co-cultures were incubated for a further 3 hours, and cells were then washed in PBS and fixed in 1% paraformaldehyde (PFA).

For all flow cytometry-based assays, cells were analysed using a Cytoflex LX (Beckman Coulter) or Attune (Life Technologies) flow cytometers and analysed using FlowJo V10.8.1 and CytExpert V2.5 software.

### ELISA

IFNα ELISA was carried out using Maxisorp plates and matched paired antibodies (Table S2). Detection of IFNβ in cell supernatants was carried out using Human IFNβ ELISA Kit (R&D Sytems).

### Transcriptome analysis

Expression of the CD99 and LDLR genes was analysed using bulk RNA sequencing (RNAseq) data from tumours and cell lines (Brohl et al., 2014, 2021) and single cell RNAseq (scRNAseq) data from tumours (Visser et al., 2023). The Coefficient of Variation (CV) of expression was calculated from the standard deviation divided by the mean of expression for each gene.

### Statistical analysis

Significant differences in results were determined using a two-tailed t-test or ANOVA with a Tukey’s post-hoc test. Correlations were determined using a Pearson’s correlation coefficient (r). IC50 values were calculated using regression analysis, applying a best fit line. Statistical analyses were performed using GraphPad PRISM 10 software.

## Results

### EWS cell lines are sensitive to MG1 infection, replication and oncolysis

Four, long-established EWS cell lines were tested for their susceptibility to direct oncolysis by MG1. SK-N-MC, SK-ES-1 and TC-32 cell lines harbour the t(11;22)(q24;q12) *EWSR1::FLI1* oncogenic fusion event, and TTC-466 the t(21;22)(q22;q12) *EWSR1::ERG* fusion oncogene. Sarcomas are derived from mesenchymal tissues and the EWS cell of origin has been reported to be mesenchymal stem cells (MSCs; also known as mesenchymal stromal cells) (Hancock & Lessnick, 2008; Miyagawa et al., 2008). We therefore compared MG1-mediated lysis of bone marrow derived MSCs from a healthy donor, with that of EWS cell lines to validate tumour selective oncolytic effects of MG1. EWS cell lines and MSCs were infected with MG1 at an increasing multiplicity of infection (MOI) and cell death was analysed 48 hours post-infection by flow cytometry. All four EWS cell lines were susceptible to MG1 infection and oncolysis, even at the lowest titre of 0.1 plaque forming units (PFU) per cell (Figure 1A; p<0.0001). By contrast, the MSCs were resistant to MG1 oncolysis, consistent with tumour selective activity of MG1 (Figure 1A).

**Figure 1:**
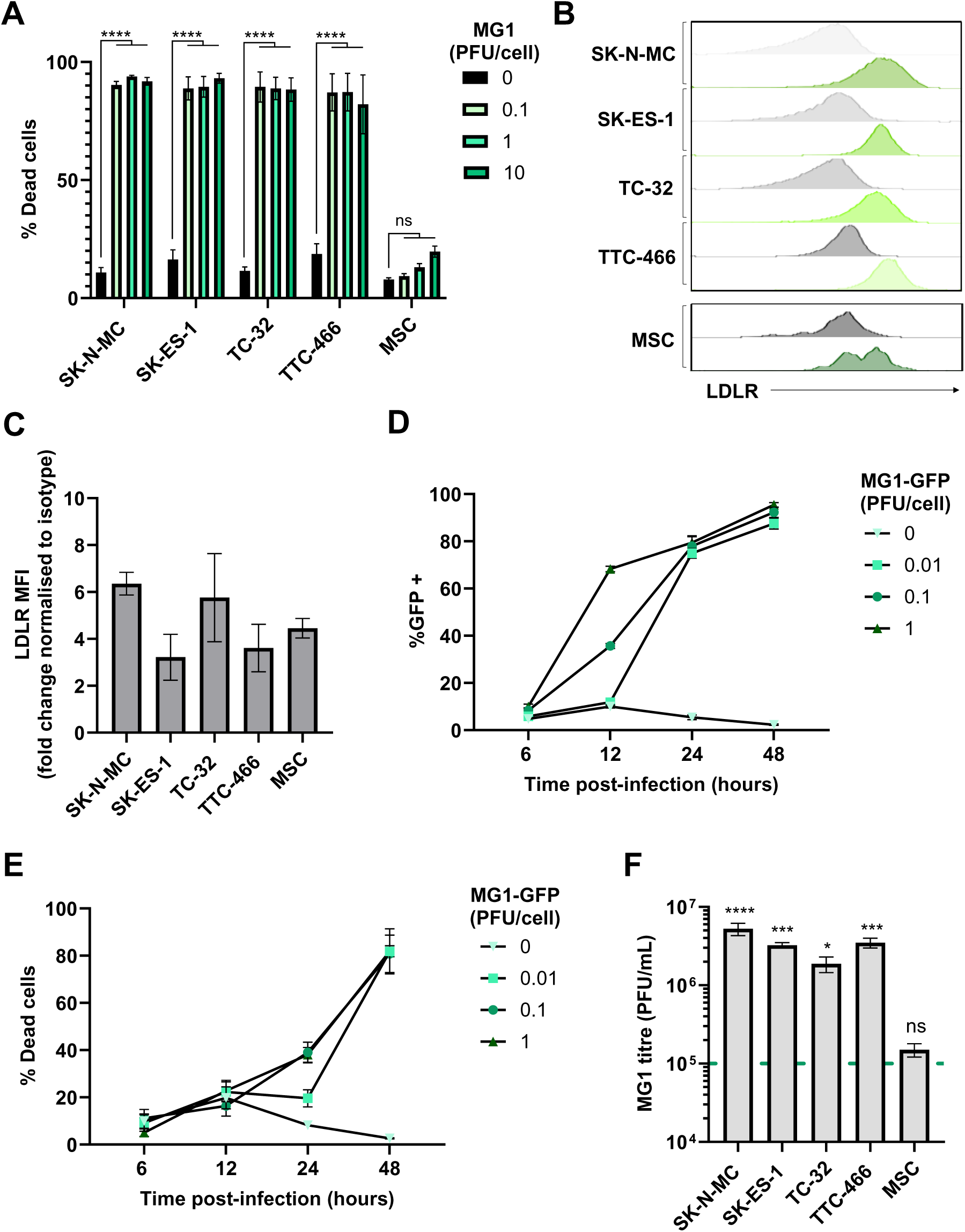
EWS cell lines are susceptible to direct oncolytic effects of MG1. (A) Four EWS cell lines (SK-N-MC, SK-ES-1, TC-32 and TTC-466) and primary healthy donor MSCs were treated ± MG1 at 0.1, 1 or 10 plaque forming units (PFU)/cell. After 48 hours, cells were stained with LIVE/DEAD™ Fixable Yellow Dead Cell Stain and analysed using flow cytometry. (B-C) EWS cell line and MSC expression of MG1 viral entry receptor LDLR was assessed using fluorescently conjugated antibodies and a matched isotype control, expression was quantified using flow cytometry, (B) grey = isotype control, green = LDLR. (C) Summary of expression of LDLR flow cytometry; the y axis shows the fold change in mean fluorescence intensity (MFI) of LDLR expression compared to staining using the isotype control antibody. (D-E) TC-32 cell lines were treated ± MG1-GFP at 0.01, 0.1, 1 PFU/cell. Cells were harvested at 6, 12, 24 and48 hours post infection and stained with LIVE/DEAD™ Fixable Yellow Dead Cell Stain. Shown are the percentage of (D) GFP positive MG1 infected cells and (E) % dead cells, which was quantified using flow cytometry. (F) EWS cell lines and MSC were treated with 1 PFU/cell MG1 for 48 hours, cell supernatants were collected and MG1 titre assessed using standard plaque assay on Vero cell line. Dashed green line represents MOI of viral input, and statistical analysis compares viral titre to viral input. All results show the mean ± standard error of the mean (SEM) for a minimum of n=3 independent experiments.

MG1 entry into cancer cells requires the ubiquitously expressed low density lipoprotein receptor (LDLR), and all the long-established cell lines and MSCs expressed cell surface LDLR as determined by flow cytometry (Figure 1B-C). We analysed the kinetics of MG1 infection of EWS using a GFP-encoding MG1 (MG1-GFP). Viral gene expression was detectable ∼12 hours post infection and cell death occurred ∼24 hours post infection (Figure 1D-E and Figure S1). Supernatants from MG1 infected cells were screened for MG1 output 48 hours post-infection, demonstrating a significant increase in viral titre for all EWS cell lines, with a greater than 10-fold increase in virus relative to the input (p<0.05; Figure 1F). Furthermore, replication was tumour selective, with no significant increase in viral titre observed following MG1 treatment of MSCs, despite their expression of LDLR, indicating while expression of this receptor may be necessary for viral infection, other cellular factors modulated in EWS cells may mediate sensitivity to replication and oncolysis (Figure 1C&F).

### Maraba virus retains oncolytic effects against chemotherapy resistant and spheroid cultured EWS cell lines

As with many other cancers, acquired resistance to chemotherapy presents a major challenge to successful EWS treatment (Stahl et al., 2011). Therefore, we generated SK-N-MC and TC-32 cell lines with resistance to doxorubicin, producing cell lines that were ∼7 times (for SK-N-MC^doxR^) and ∼24 times (for TC-32^doxR^) more resistant to doxorubicin chemotherapy than their parental counterparts, based on IC_50_ values (Figure 2A-B). Importantly, these doxorubicin resistant derivatives did not display any significant differences in cell surface LDLR expression or sensitivity to MG1 oncolysis (Figure 2C-D).

**Figure 2:**
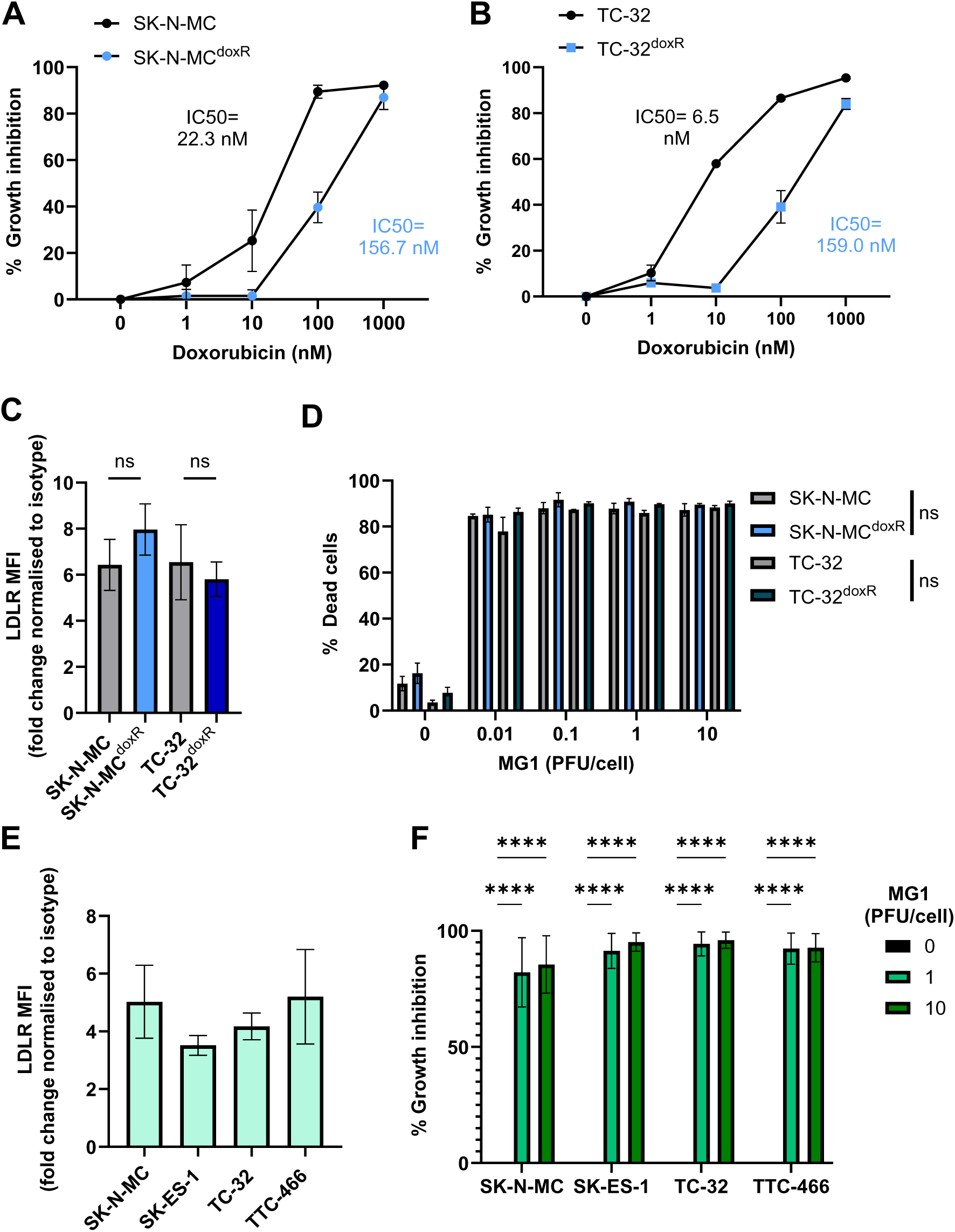
Doxorubicin resistant EWS cell lines and spheroid cultures retain sensitivity to MG1 oncolysis. (A-B) Doxorubicin-resistant (SK-N-MC^doxR^ and TC-32^doxR^) and parental (SK-N-MC and TC-32) cell lines were treated with doxorubicin concentrations ranging from 0-1000 nM for 48 hours. Growth inhibition was assessed using an MTT assay and the IC50 values calculated using nonlinear regression analysis. (C) Expression of MG1 entry receptor LDLR on doxorubicin-resistant and parental cell lines was assessed using flow cytometry, results show the fold change in MFI relative to isotype controls. (D) Doxorubicin-resistant and parental cell lines were treated ± MG1 at 0.01, 0.1, 1 and 10 PFU/cell for 48 hours. Cells were stained with LIVE/DEAD™ Fixable Yellow Dead Cell Stain and cell death determined using flow cytometry. (E) Expression of LDLR on EWS spheroid cultures was assessed by dissociation of spheroids using Accutase. Dissociated cells were stained with LIVE/DEAD™ Fixable Yellow Dead Cell Stain and fluorescently conjugated anti-LDLR antibodies, and a matched isotype control, results show the fold change in MFI on live cells, relative to isotype controls. (F) EWS spheroids were treated ± MG1 at 1 or 10 PFU/cell for 48 hours and CellTiter-Glo® reagent was used to measure growth inhibition. Results were normalised to an untreated control. All results show the mean ± SEM for a minimum of n=3 independent experiments.

Spheroid cultures more closely mimic solid tumour architecture than monolayer cultures, and demonstrate greater resistance to certain therapeutics, for example showing reduced susceptibility to chemotherapy (Fong et al., 2013). Moreover, MG1 cytotoxicity has been reported to be reduced in ovarian cancer spheroids, as a result of reduced LDLR expression, therefore we wanted to investigate if similar effects were seen in EWS (Tong et al., 2015). We generated spheroids from the four long-established cell lines (Figure S2). Unlike results from ovarian cancer, the EWS cell spheroids expressed LDLR and remained susceptible to MG1 oncolysis (Figure 2E-F). These results demonstrate that MG1 infects and kills EWS in both conventional 2D culture conditions and when grown as 3D spheroids.

### PDES cell cultures express LDLR and retain sensitivity to MG1 oncolysis

Next, we investigated the ability of MG1 to infect and kill PDES cell cultures. PDES cultures were more recently established from patient samples and for these studies we used three PDES cell cultures, two of which express EWSR1::FLI1 fusion oncoprotein (CCRG1-L-017 and CCRG-L-023) and one which expresses EWSR1::ERG fusion oncoprotein (CCRG1-L-066) (Roundhill et al., 2021). All of the PDES cell cultures express cell surface CD99 (a protein over-expressed in EWS tumours and used in diagnosis), albeit with greater heterogeneity than the established cell lines (Figure S3A). The PDES cells are transcriptionally distinct from long-established EWS cell lines and are more resistant to chemotherapy (Roundhill et al., 2021, 2023). Furthermore, PDES cell cultures have a different forward and side scatter profile when assessed by flow cytometry and microscopy demonstrating that they are larger with a more elongated, mesenchymal-like morphology (Figure S3B-C). Importantly, all three PDES cell cultures demonstrated susceptibility to MG1, which was consistent with LDLR expression (Figure 3A-B).

**Figure 3:**
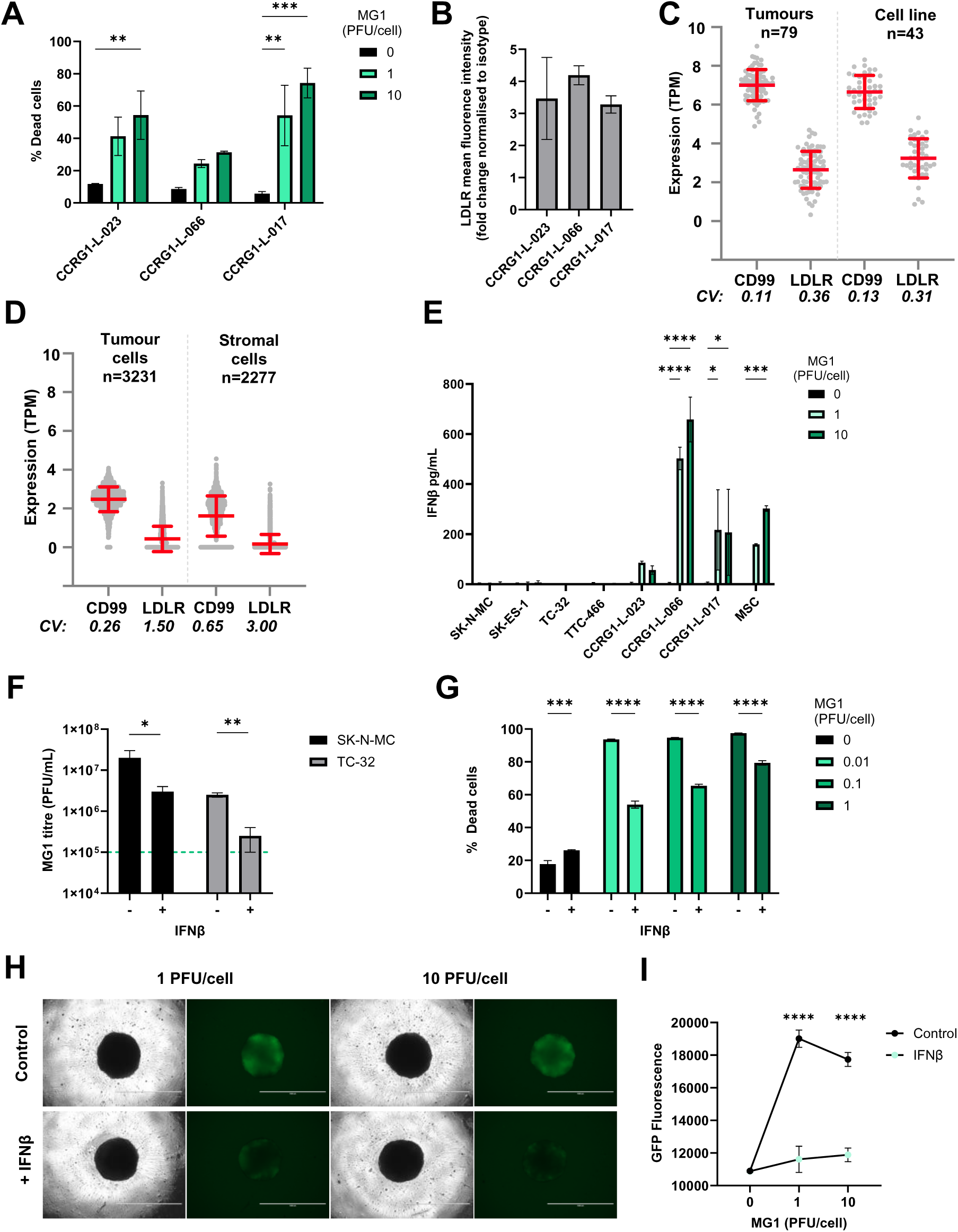
Sensitivity of EWS to MG1 oncolysis is modulated by IFNβ. (A) PDES cell cultures were treated ± MG1 at 1 or 10 PFU/cell for 48 hours. Cells were stained with LIVE/DEAD™ Fixable Yellow Dead Cell Stain and cell death assessed using flow cytometry. (B) PDES cell culture expression of viral entry receptor LDLR assessed using flow cytometry, results show the fold change in MFI relative to isotype controls. (C-D) Expression of CD99 and LDLR genes was analysed using RNA sequencing data, (C) bulk RNAseq data from 43 EWS cell lines and 79 EWS tumour samples (Brohl et al., 2014, 2021) and (D) single cell RNA sequencing data from 3231 tumour cells and 2277 stromal cells (Visser et al., 2023). The Coefficient of Variation (CV) of expression was calculated from the standard deviation divided by the mean of expression for each gene. (E) PDES cell cultures and established cell lines were treated ± MG1 at 1 or 10 PFU/cell and cell-free supernatants were collected after 48 hours and screened for IFNβ using ELISA. (F) TC-32 and SK-N-MC cells were treated ± IFNβ at 400 pg/mL for 24 hours and then treated ± MG1 at 1 PFU/cell for 24 hours. Cell free supernatants were screened for MG1 using plaque assay to assess viral replication, the green dashed line represents titre of viral input. (G) TC-32 cells were treated ± IFNβ at 400 pg/mL for 24 hours, and then treated ± MG1 at 0.01, 0.1 and 1 PFU/cell for 48 hours. Cells were stained with LIVE/DEAD™ Fixable Yellow Dead Cell Stain and analysed using flow cytometry. (H-I) TC-32 cells were seeded into low adhesion 96 well plates at 2000 cells/well to generate spheroids over 7 days. Spheroids were treated ± IFNβ at 400 pg/mL for 24 hours and then treated ± MG1-GFP at 1 or 10 PFU/cell. After 24 hours, (H) cells were imaged to assess GFP fluorescence using EVOS microscope at 4x magnification, scale bar represents 1 mm. (I) GFP fluorescence was quantified using Cytation 5 plate reader. (A-B; E-I) results presented the mean ± SEM for a minimum of n=3 independent experiments.

To determine the potential utility of MG1 as a therapeutic agent for EWS patients, we next evaluated LDLR and CD99 expression in a larger cohort of EWS samples using transcriptome data. CD99 is highly expressed on EWS cells, where it serves as a key diagnostic marker. However, its expression is not limited to EWS; it is also found on a variety of other cell types including immune cells, mesenchymal cells, and endothelial cells (Pasello et al., 2018). A cohort of 79 EWS tumours and 43 EWS cell lines (Brohl et al., 2021) showed that CD99 and LDLR gene expression was detected in all of the patient samples and established cell lines tested (Figure 3C). Furthermore, to address heterogeneity at the single cell level we analysed EWS scRNAseq data (Visser et al., 2023). Within this dataset, Visser *et al* identified both tumour cells (n=3231) and stromal cells (n=2277), the latter comprising mesenchymal cells, endothelial cells, and multiple immune cell types. Expression of both CD99 and LDLR genes was variable in the stroma, reflecting the multiple cell types present in this compartment (Figure 3D). As expected, CD99 expression was relatively uniform in the EWS tumour cells (98% of tumour cells expressed CD99), but only 38% of EWS tumour cells expressed detectable levels of LDLR mRNA (Figure 3D). Heterogeneity of LDLR gene expression at the single cell level suggests that not all malignant cells within the EWS tumour will be directly targetable by MG1 infection and oncolysis.

### EWS sensitivity to Maraba virus is counteracted by IFNβ responses

The long established EWS cell lines and PDES cultures express LDLR but displayed variable susceptibility to MG1. For example, treatment with 1 PFU/cell MG1 was sufficient to kill ∼80-90% of the population of the established cell lines (Figure 1A), whereas the same titre killed less than 60% of the population in PDES cell cultures (Figure 3A). We speculated, given the known IFN-I sensitive nature of MG1 previously reported, that the PDES cell cultures might retain some anti-viral activity and thereby restrict MG1 replication (Brun et al., 2010). In support of this, all three PDES cultures (and the MSC culture) produced detectable levels of IFNβ in response to MG1 infection, whereas no IFNβ was detected upon infection of the established cell lines (Figure 3E). None of the cell types tested secreted IFNα in response to MG1 (Figure S4A).

The importance of IFNβ in determining the susceptibility to MG1 was further investigated by performing infections in the presence or absence of exogenous IFNβ; TC-32 and SK-N-MC cell lines were pre-treated with recombinant human IFNβ for 24 hours before MG1 infection and tested for the effect of this IFN-I on viral replication and oncolysis. These data show that IFNβ treatment significantly reduced MG1 replication and oncolysis in established EWS cell lines (Figure 3F-G and Figure S4B). This effect was also observed in spheroids, where IFNβ pre-treatment of TC-32 and SK-N-MC spheroid cultures significantly reduced MG1-GFP replication (Figure 3H-I and S4C-D). These results demonstrate that the anti-viral capacity of the EWS tumour cells is an important determinant of their susceptibility to MG1.

### MG1 activates NK cells and granule mediated destruction of EWS

Although IFN-I can limit viral replication, its production is critical for triggering host anti-tumour immune responses, including NK cell activation and cytotoxicity (Marelli et al., 2018; Wantoch et al., 2022). To test this possibility in EWS, we treated PBMC isolated from healthy donors with an increasing MOI of MG1 for 48 hours. As expected, this resulted in production of IFN-I (IFNα and IFNβ), and expression of the IFN-I inducible molecules CD69 and CD317/tetherin on the NK cell surface (Figure 4A-C). To examine NK cell cytotoxicity, we repeated treatment of PBMC with MG1 (at 1 PFU/cell) for 48 hours and performed an immune cytotoxicity assay by co-culturing the virus-stimulated PBMC with EWS target cells for 5 hours and assaying EWS cell death (Figure 4D). Importantly, MG1 does not kill EWS cell lines within 5 hours (as shown in Figure 1A) and hence this assay reflects PBMC-mediated killing and not virus-mediated oncolysis; PBMC-mediated killing of SK-N-MC, SK-ES-1, TC-32 and TTC-466 cells was significantly enhanced following pre-treatment of PBMC with MG1 (Figure 4E). Furthermore, doxorubicin resistant cell cultures SK-N-MC^doxR^ and TC-32^doxR^ and PDES cell cultures were also sensitive to the action of MG1 treated PBMC (Figure 4F-G).

**Figure 4:**
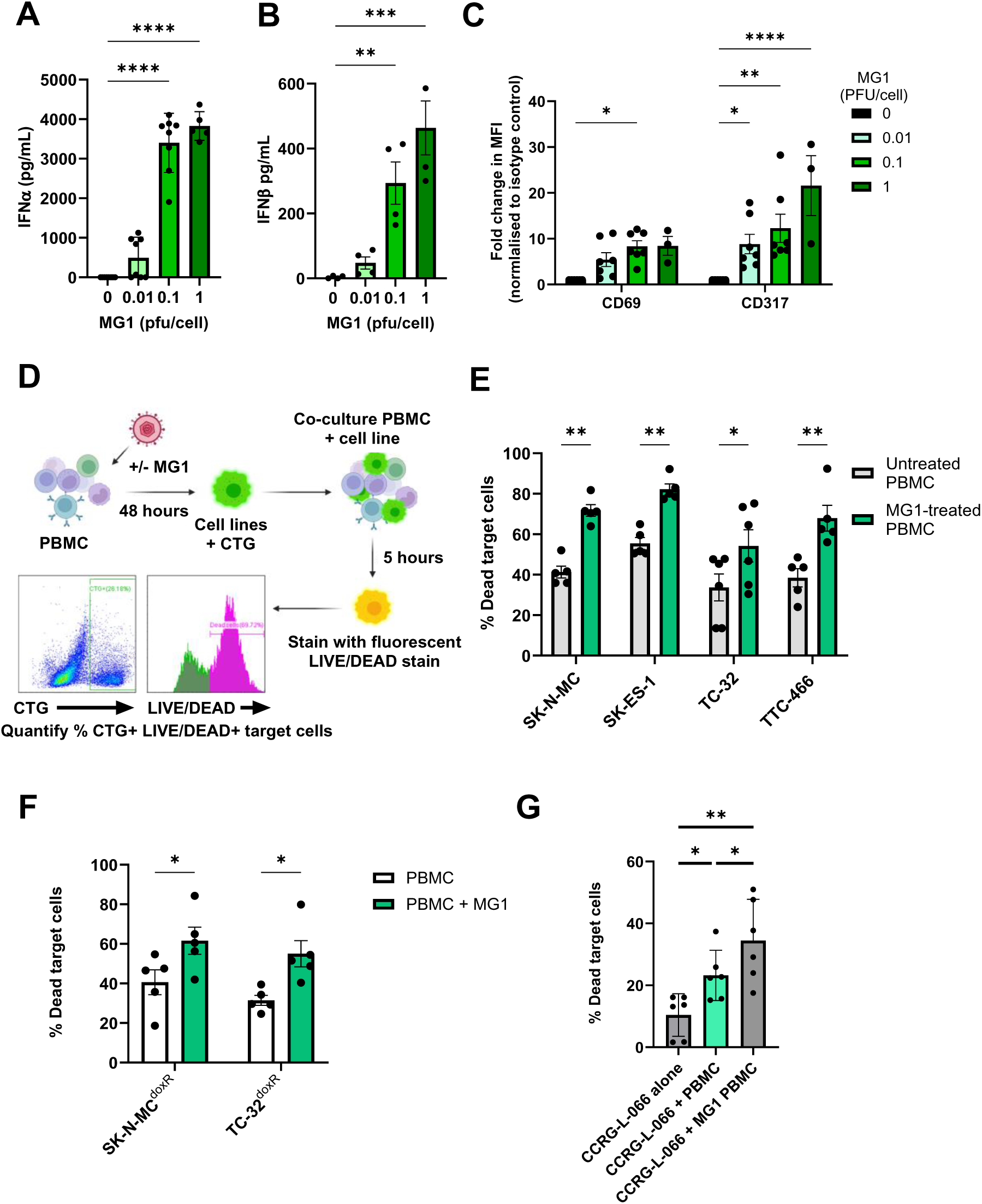
MG1 treatment stimulates immune mediated killing of EWS. (A-C) PBMC were treated ± MG1 at 0.01, 0.1 or 1 PFU/cell for 48 hours. (A-B) Cell-free supernatants were collected and (A) IFNα and (B) IFNβ detected using ELISA. (C) Cells were stained with CD56, CD3, CD69 or CD317 antibodies, or matched isotype controls. Cells were analysed to assess MFI of CD69 and CD317 activation markers on NK cells (CD56+CD3-) using flow cytometry. Schematic showing process of immune killing assays. Briefly, PBMC were treated ± 1 PFU/cell MG1 for 48 hours and then co-cultured with cell tracker green (CTG) labelled EWS cells for 5 hours. Cells were then stained with LIVE/DEAD™ Fixable Yellow Dead Cell Stain, and the percentage of dead target cells quantified by flow cytometry. Immune killing assay of EWS cell lines, (F) doxorubicin-resistant cell lines and (G) patient-derived cell culture CCRG-L-066. All results show the mean ± SEM for a minimum of n=3 independent experiments.

To investigate the importance of NK cells in this activity, we repeated these experiments using PBMC from which the NK cells had been depleted using magnetic immunoselection. Flow cytometry confirmed successful depletion of the NK cells from PBMC (∼86% depleted, Figure 5A), which resulted in a significant reduction in killing of the four EWS cell lines (Figure 5B-C and Figure S5A-B). These results demonstrate that MG1 activates NK cell-mediated cytotoxicity towards EWS cell lines. Accordingly, NK cell degranulation in response to EWS cell lines (cultured as either monolayers or spheroids) was significantly enhanced by pre-treatment with MG1 (Figure 5D-E and Figure S5C), suggesting that perforin/granzyme mediated cytotoxicity was responsible for EWS cell death. Delivery of granzymes to target cells is dependent on calcium-mediated oligomerisation of perforin, and treatment of PBMC with the calcium chelator EGTA inhibited unstimulated and MG1 stimulated PBMC-mediated killing of both SK-N-MC and TTC-466 EWS cell lines, supporting a role for perforin-dependent granule-mediated killing of the EWS cells (Figure 5F and Figure S5D) (Lopez et al., 2013).

**Figure 5:**
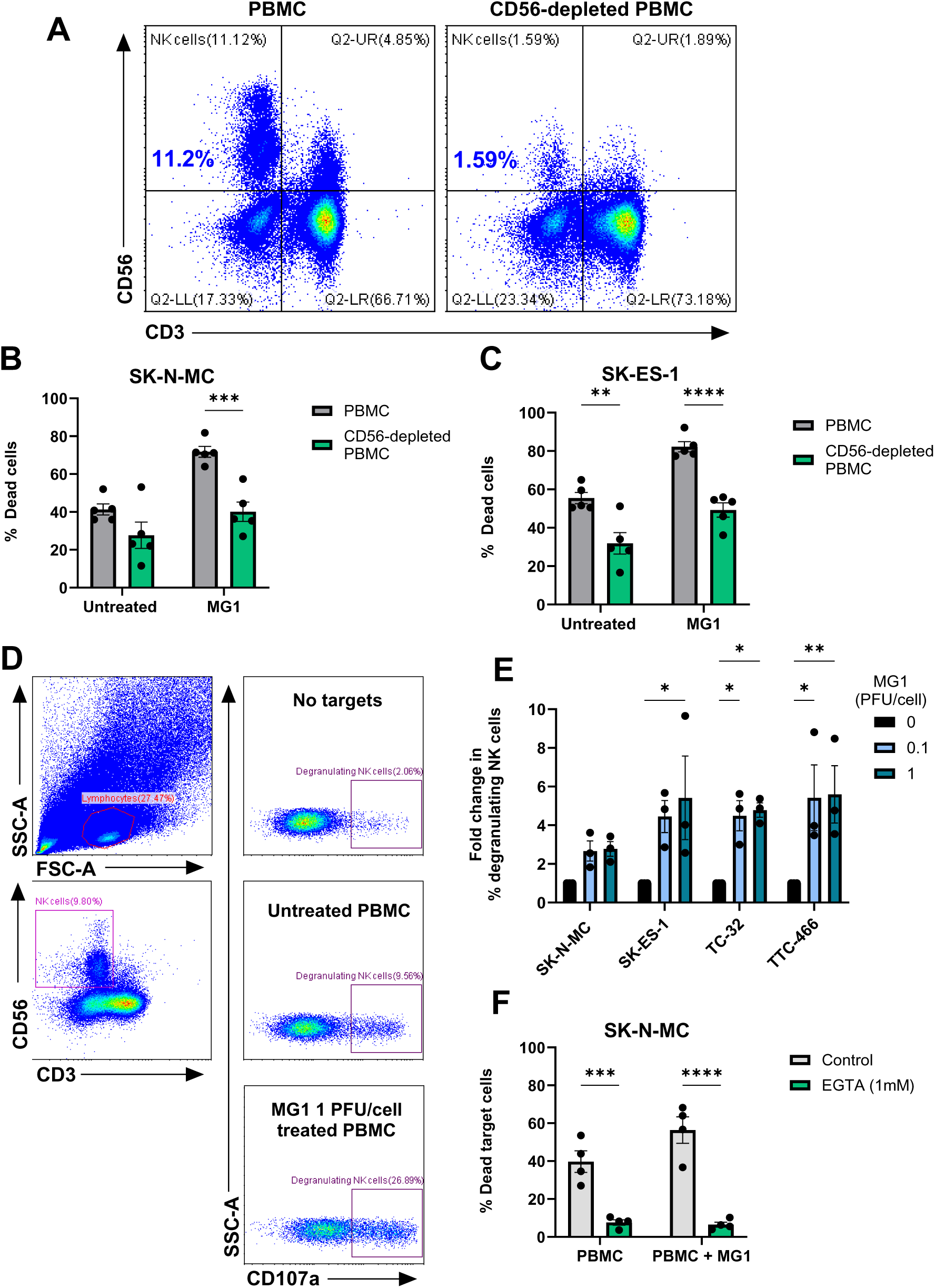
MG1 stimulates immune-mediated killing of EWS targets by NK degranulation. (A-C) NK cells were depleted from PBMC using CD56 magnetic bead selection and PBMC were treated ± MG1 at 1 PFU/cell for 48 hours. (A) Depletion was validated using anti-CD56 and anti-CD3 antibodies and flow cytometry. (B-C) Whole PBMC or CD56-depleted PBMC were co-cultured with cell tracker green stained (B) SK-N-MC, (C) SK-ES-1 EWS target cells at a ratio of 25:1 for 5 hours. Co-cultures were stained with LIVE/DEAD™ Fixable Yellow Dead Cell Stain and the percentage of dead target cells assessed by flow cytometry. (D-E) PBMC were treated ± MG1 at 1 PFU/cell for 48 hours. PBMC were co-cultured at a ratio of 10:1 with EWS cell lines for 4 hours. The percentage of degranulating NK cells was detected by staining with CD56, CD3 and CD107a antibodies and flow cytometry. (D) Representative gating strategy for SK-N-MC target cells, (E) summary of results presented as fold change in degranulating NK cells relative to untreated PBMC control. (F) PBMC were treated ± MG1 at 1 PFU/cell for 48 hours and then treated ±1 mM EGTA for 30 minutes. PBMC were then co-cultured with SK-N-MC EWS target cells at a ratio of 25:1 for 5 hours. Cells were stained with LIVE/DEAD™ Fixable Yellow Dead Cell Stain and the percentage dead target cells assessed using flow cytometry. All results show the mean ± SEM for a minimum of n=3 independent experiments.

## Discussion

Oncolytic viruses offer an innovative approach for stimulating the immune response against hard-to-treat solid tumours such as EWS. In this study, we present a comprehensive study of the effect of MG1 against EWS, and demonstrate that the oncolytic rhabdovirus, Maraba virus MG1, is a candidate OV for the treatment of EWS. To highlight the potential clinical relevance of our study, we used complementary EWS model systems, including those recently derived from patient samples and cell lines that exhibit resistance to drugs used as part of the standard - of-care for EWS. Mechanistic studies show that MG1 can kill EWS cells directly and mediate immune-based killing of EWS cells, specifically through the action of NK cells.

When comparing OV, MG1 frequently out-performs other rhabdoviruses, and OV from other virus families in terms of direct oncolytic action *in vitro* (Brun et al., 2010; Le Boeuf et al., 2017; Tong et al., 2015). Previous publications have demonstrated that MG1 induces direct lysis of several solid cancer cell types including melanoma, breast, colon, ovarian and lung cancer cell lines (Alkayyal et al., 2017; Armstrong et al., 2024; M.-C. Bourgeois-Daigneault et al., 2018; M. C. Bourgeois-Daigneault et al., 2016; Brun et al., 2010; Mahoney et al., 2011). However, the action of MG1 against EWS remains relatively unexplored. Le Bouef *et al* demonstrated direct oncolytic activity of MG1 against the long established EWS cell line, RD-ES, in monolayer cultures. The same study showed that primary tumour tissue isolated from a variety of sarcoma types could support MG1 replication and that murine S180 sarcomas (of fibrosarcoma origin) were susceptible to MG1 *in vivo* (Le Boeuf et al., 2017). However, these studies did not include EWS patient samples or EWS cell lines other than RD-ES. Our findings build substantially on this previous work, establishing the efficacy of MG1 in more complex 3D spheroid models, chemotherapy-resistant cell lines and PDES cell cultures and demonstrate that MG1 is capable of infecting, replicating, and lysing clinically relevant EWS cell models *in vitro*.

Studies of MG1 and ovarian cancer demonstrated that cell surface expression of LDLR was required for MG1 entry and that LDLR expression was reduced when culturing cells as spheroids compared to conventional monolayers, delaying oncolysis (Tong et al., 2015). By contrast, our data demonstrated that LDLR was expressed in all the EWS cell models tested, and all were susceptible to MG1 oncolysis, albeit to varying levels. However, transcriptome data demonstrated heterogeneity of LDLR gene expression at the single cell level, with substantial populations of EWS tumour cells lacking detectable LDLR gene expression. Heterogeneity of LDLR expression could be viewed as a barrier to the use of MG1 in EWS, as large populations of tumour cells may not be amenable to infection. However, LDLR expression is dynamic and the *LDLR* gene, along with genes encoding components of cholesterol biosynthesis, are expressed in response to a drop in cellular cholesterol. This allows cells to endocytose cholesterol from the extracellular environment (via LDLR) and to synthesise new cholesterol, restoring the pool (Horton et al., 2002). The heterogeneity of *LDLR* gene expression observed in EWS scRNAseq might therefore reflect a snapshot of the differential requirements for cholesterol amongst tumour cells, for example a greater requirement in those cells undergoing proliferation at the time of analysis.

Resistance to oncolysis affects just one arm of the action of OV. Building on models of OV action, we suggest that there will be oncolysis of some EWS cells and this will release damage associated molecular patterns, pathogen associated molecular patterns and tumour antigens (Lin et al., 2023). This will occur within the context of an anti-viral response mediated by non-malignant cells, including the production of IFN-I and the activation of NK cells. We have previously shown that OV can stimulate the production of IFN-I from monocytes, that the activation of NK cells by OV is IFN-I dependent *in vitro* and that the peak of IFN-I responses coincides with the peak of NK cell activation in patients receiving intravenous OV (Parrish et al., 2015; Wantoch et al., 2022). Furthermore, OV treatment of PBMC results in the induction of expression of NK cell cytotoxic machinery, enhancing the ability of OV activated NK cells to kill tumour cells. Populations of EWS cells that are not infected and lysed by MG1 directly could nevertheless find themselves in an environment in which NK cells are activated and capable of EWS recognition and killing. Whilst IFN-I production by healthy cells will impair EWS oncolysis, it is expected to promote NK cell activation and tumour lysis. Further, although not tested here, many studies have demonstrated that OV promote T cell activity against tumours, for example, MG1 monotherapy enhanced T cell responses in melanoma models (Armstrong et al., 2024).

Maraba virus is a (+)ssRNA virus amenable to genetic manipulation and its activity can be modulated by the inclusion of transgenes. For example, MG1 encoding the cancer/testis antigen MAGE-A3 has been used as part of an OV/tumour vaccine approach and MG1 encoding microRNA molecules have been used to skew tumour infiltrating macrophages from an M2-like (pro-tumour) immunosuppressive phenotype to an M1-like pro-inflammatory/anti-tumour phenotype in ovarian cancer models (Jennings et al., 2024). Similar modifications may have the potential to enhance the activity of MG1 against EWS. Furthermore, combining the action of OV with other immunotherapeutic approaches, such as immune checkpoint inhibitors or chimeric antigen receptor (CAR)-T cells is gaining traction in the treatment of other cancer types (Lin et al., 2023).

To date, OV have demonstrated a good safety record in clinical trials (Gao et al., 2021; Schirrmacher, 2020). Restriction of OV activity by IFN-I greatly reduces the infection of healthy tissue. Not surprisingly, side effects associated with OV frequently resemble the symptoms of acute viral infection and are usually mild and transient in nature. A wild type Maraba virus strain has been shown to cause neuropathology in mice, but studies of MG1-based OV in cats and non-human primates both reported an absence of adverse effects requiring clinical intervention (Hummel et al., 2017; Maia-Farias et al., 2020; Pol et al., 2019). Furthermore, several clinical trials incorporating MG1 are underway (Clinical Trial Identifiers; NCT02285816, NCT02879760, NCT03618953, NCT03773744) and one preliminary trial report suggests that adverse effects are similar to those observed with other OV (Jonker et al., 2017). Although these clinical trials investigating MG1-based therapies for adult solid tumours are restricted to patients over 18 years of age, a separate phase I study evaluating a herpes simplex virus-based OV in paediatric and young adult patients (aged 12 to 21), including patients with bone and soft tissue sarcomas, demonstrated safety profiles comparable to those seen in adult populations (Moreno et al., 2023).

In summary, using a variety of EWS cell models, we have shown that the OV Maraba virus MG1 can infect, replicate, and kill EWS cells directly and activate NK cell cytotoxicity against this tumour. This dual activity suggests that MG1 and engineered derivatives hold promise in the future development of EWS immunotherapy.

---

**Figure S1:**
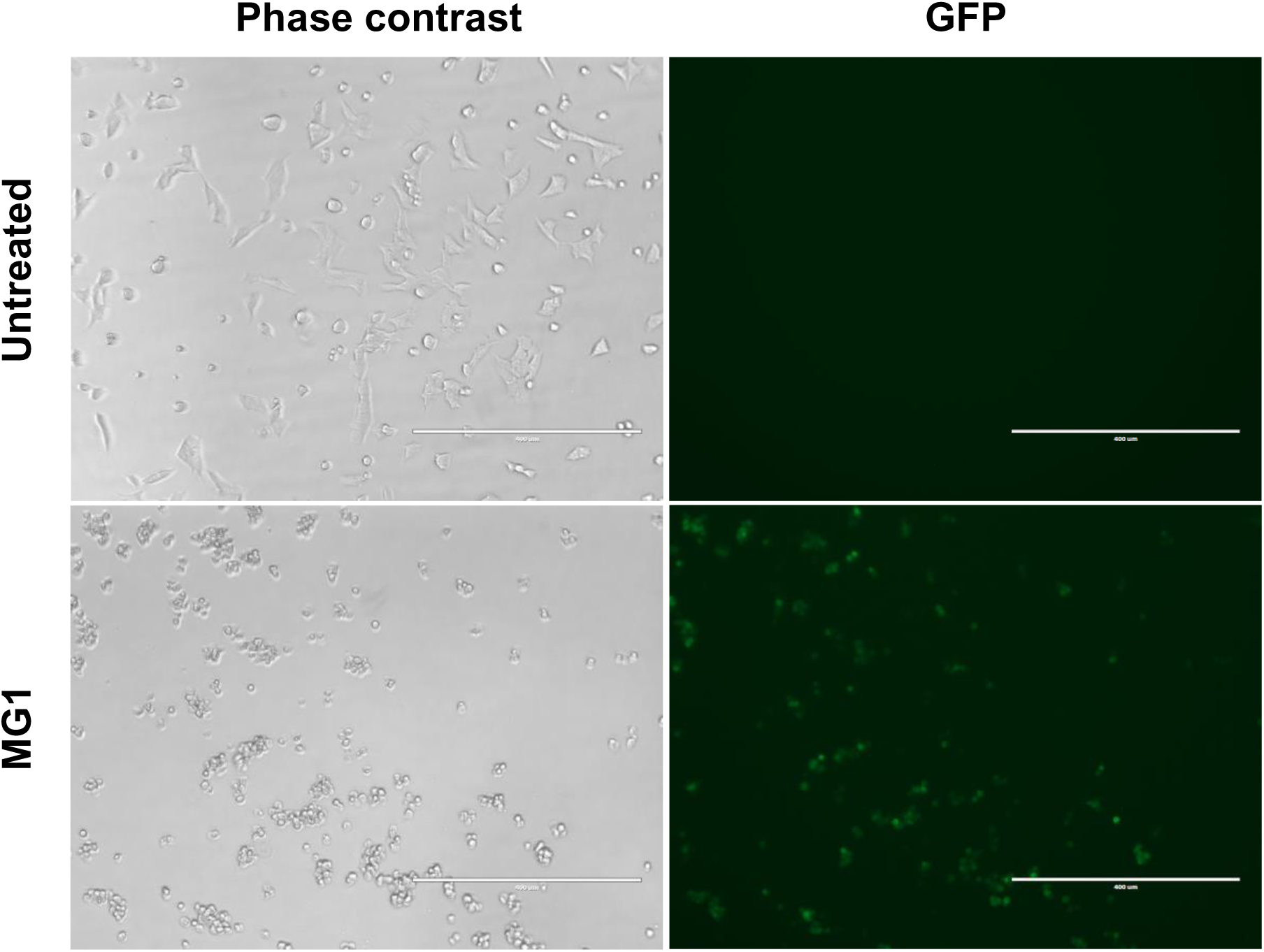
EWS cell line MG1-GFP infection. TC-32 cell lines were treated ± MG1-GFP at 1 PFU/cell. After 24 hours cells were imaged using EVOS fluorescent microscope and phase contrast and GFP images taken at 10x magnification. Scale bar = 400 µm.

**Figure S2:**
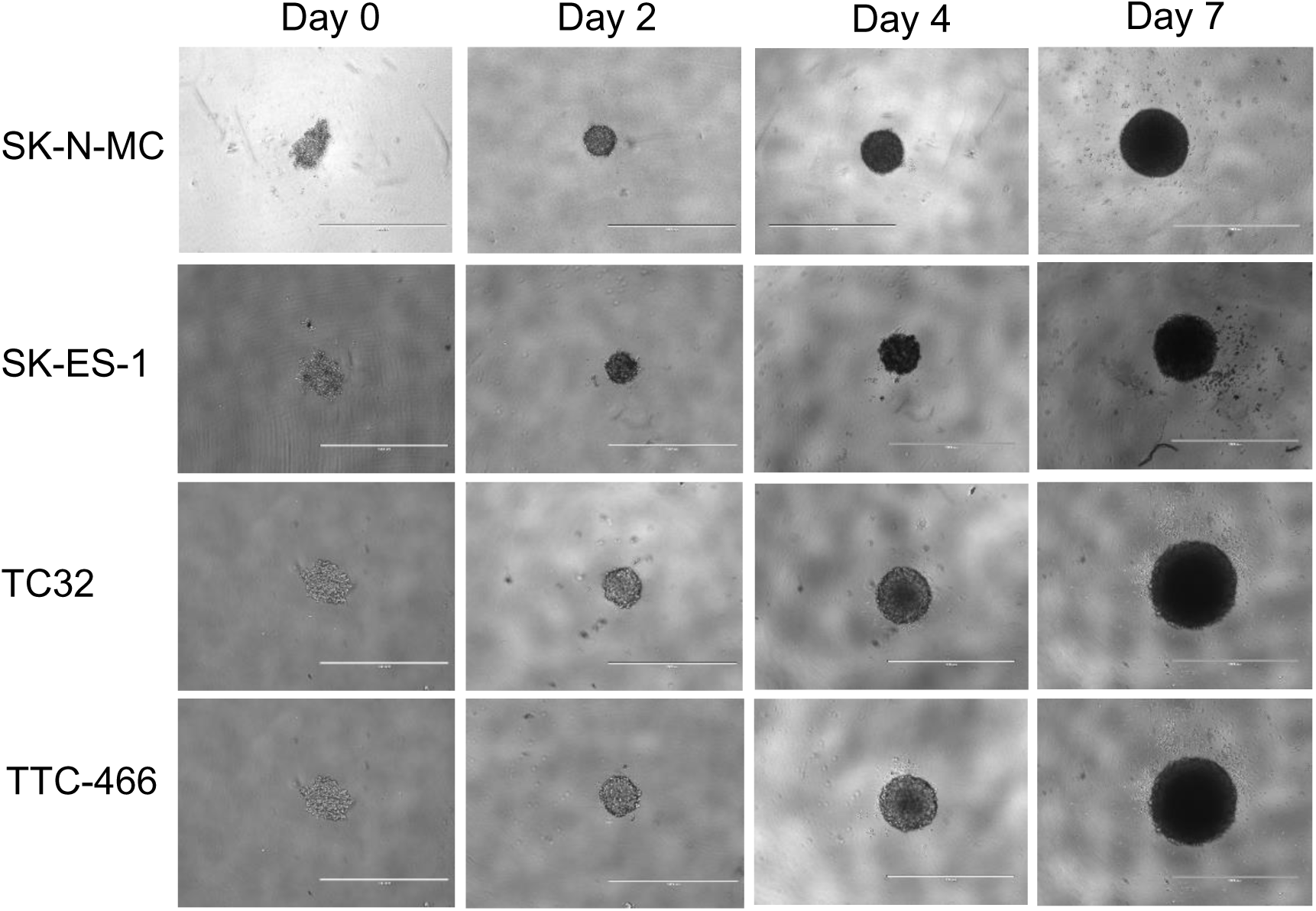
EWS cell line spheroid cultures. EWS cell lines were seeded into low adhesion 96 well plates at 2000 cells/ well for 7 days. Spheroid formation was tracked over 7 days, where spheroids were imaged using EVOS microscope at 4x magnification on days 0, 2, 4 and 7. Scale bar = 1 mm.

**Figure S3:**
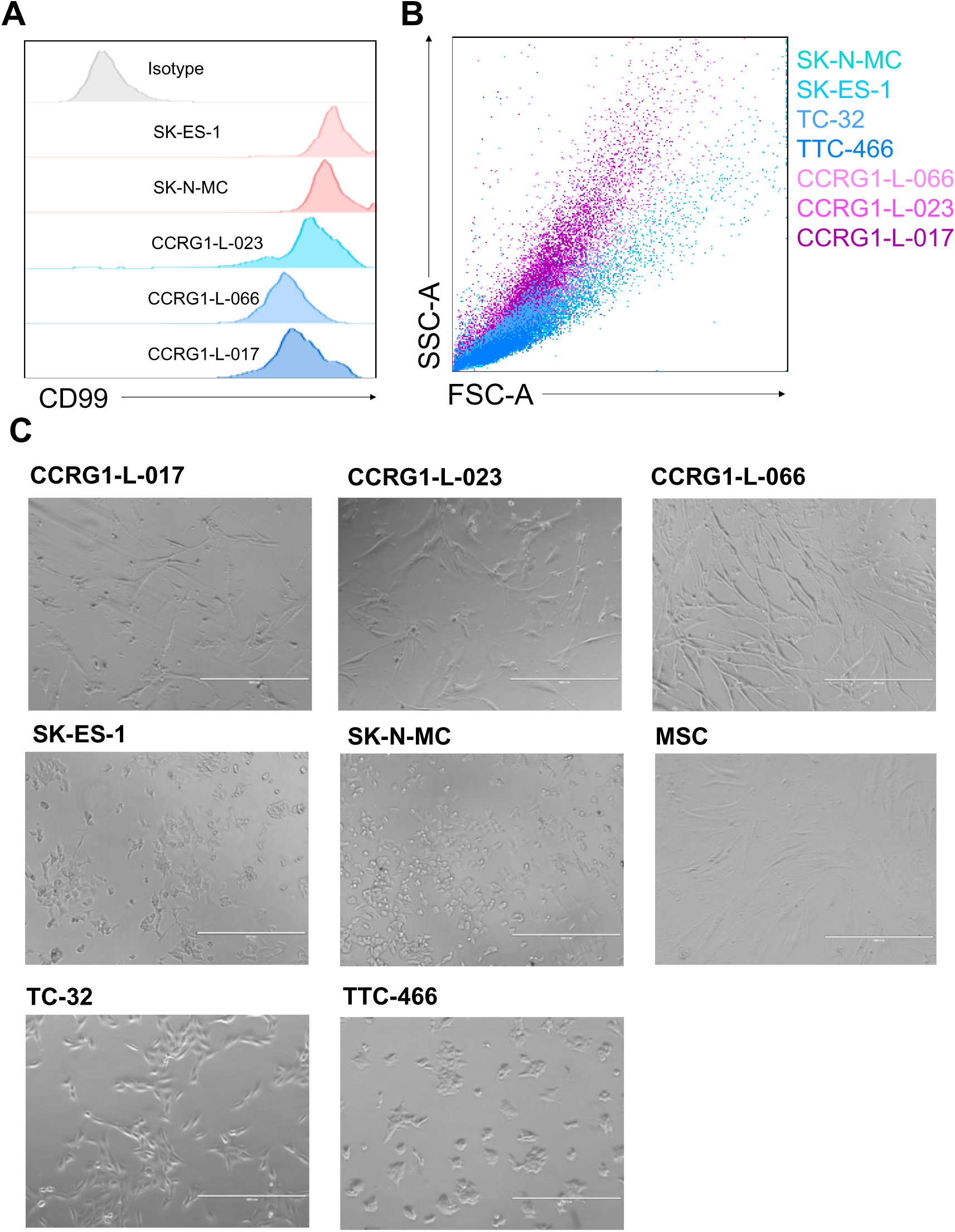
Characterisation of EWS cell cultures. (A) EWS cell lines (SK-ES-1 and SK-N-MC) and PDES cell cultures (CCRG-L-023, CCRG-L-066, CCRG-L-017) were stained with a fluorescently conjugated anti-CD99 antibody or a matched isotype control, and CD99 expression assessed using flow cytometry. (B) PDES cultures (pink) and EWS cell lines (blue) FCS-SSC profiles assessed using flow cytometry, and overlayed using FlowJo V10.8.1 software. (C) PDES cell cultures, EWS cell lines and MSC were seeded into 12 well plates at 1×10^5^ cells/well for 24 hours. Cell cultures were imaged at 10x magnification. Scale bar = 400 µm.

**Figure S4:**
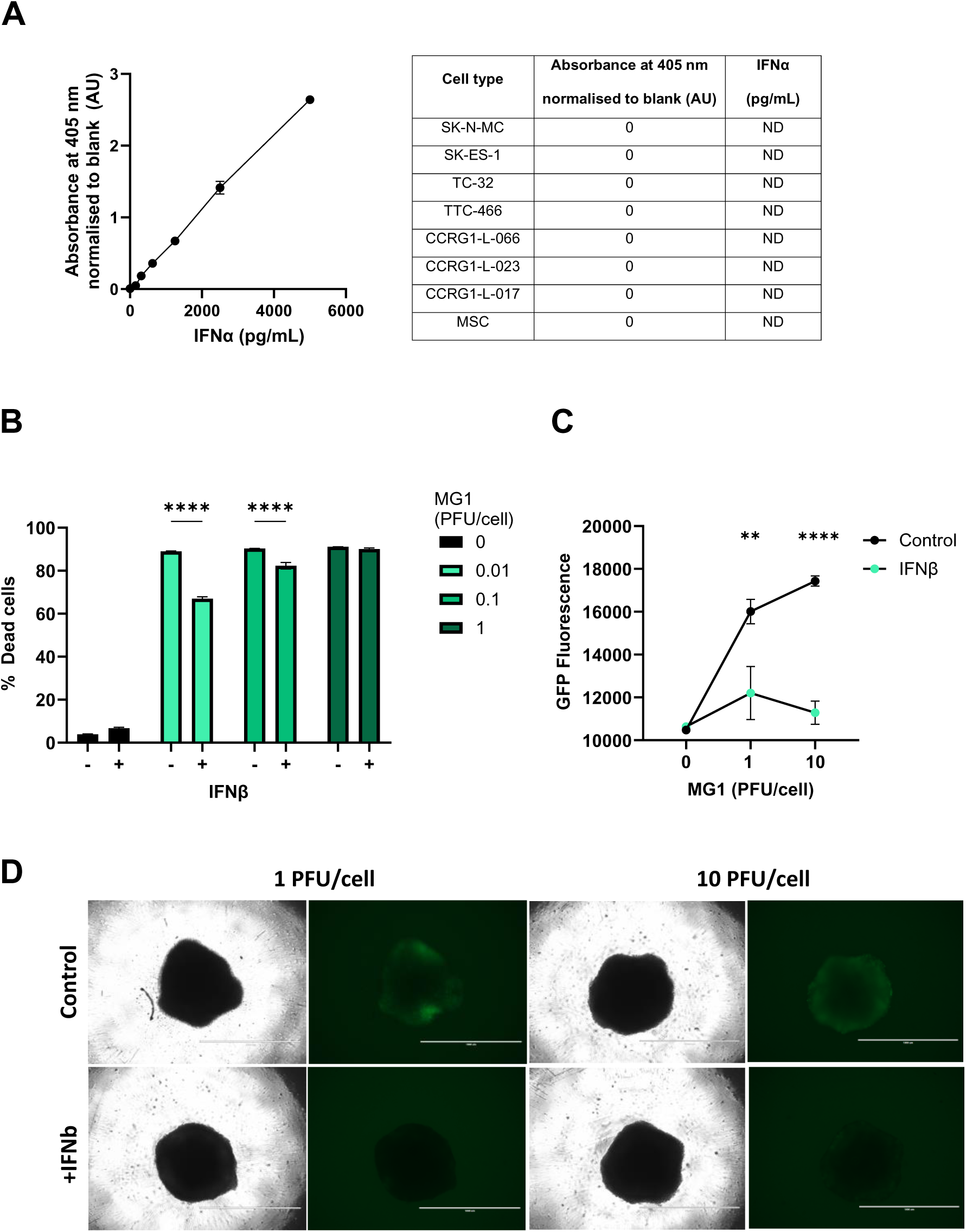
IFNβ protects EWS cell lines against MG1 replication. (A) PDES cell cultures, established cell lines and MSC were treated ± MG1 at 1 PFU/cell and cell-free supernatants were collected after 48 hours and screened alongside recombinant standards for IFNα using ELISA. Results show standard curve for ELISA standard and outputs for cell supernatants, ND = not detected. (B) SK-N-MC cells were treated ± IFNβ at 400 pg/mL for 24 hours, and then treated ± MG1 at 0.01, 0.1 and 1 PFU/cell for 48 hours. Cells were stained with LIVE/DEAD™ Fixable Yellow Dead Cell Stain and analysed using flow cytometry. (C-D) SK-N-MC cells were seeded into low adhesion 96 well plates at 2000 cells/well to generate spheroids over 7 days. Spheroids were treated ± IFNβ at 400 pg/mL for 24 hours and then treated ± MG1-GFP at 1 or 10 PFU/cell. After 24 hours (C) GFP fluorescence was quantified using Cytation 5 plate reader at wavelength 485nm and (D) cells were imaged using EVOS fluorescent microscope to obtain phase contrast and GFP images taken at 4x magnification. Scale bar = 1mm. Results presented are representative of a minimum of n=3 independent experiments.

**Figure S5:**
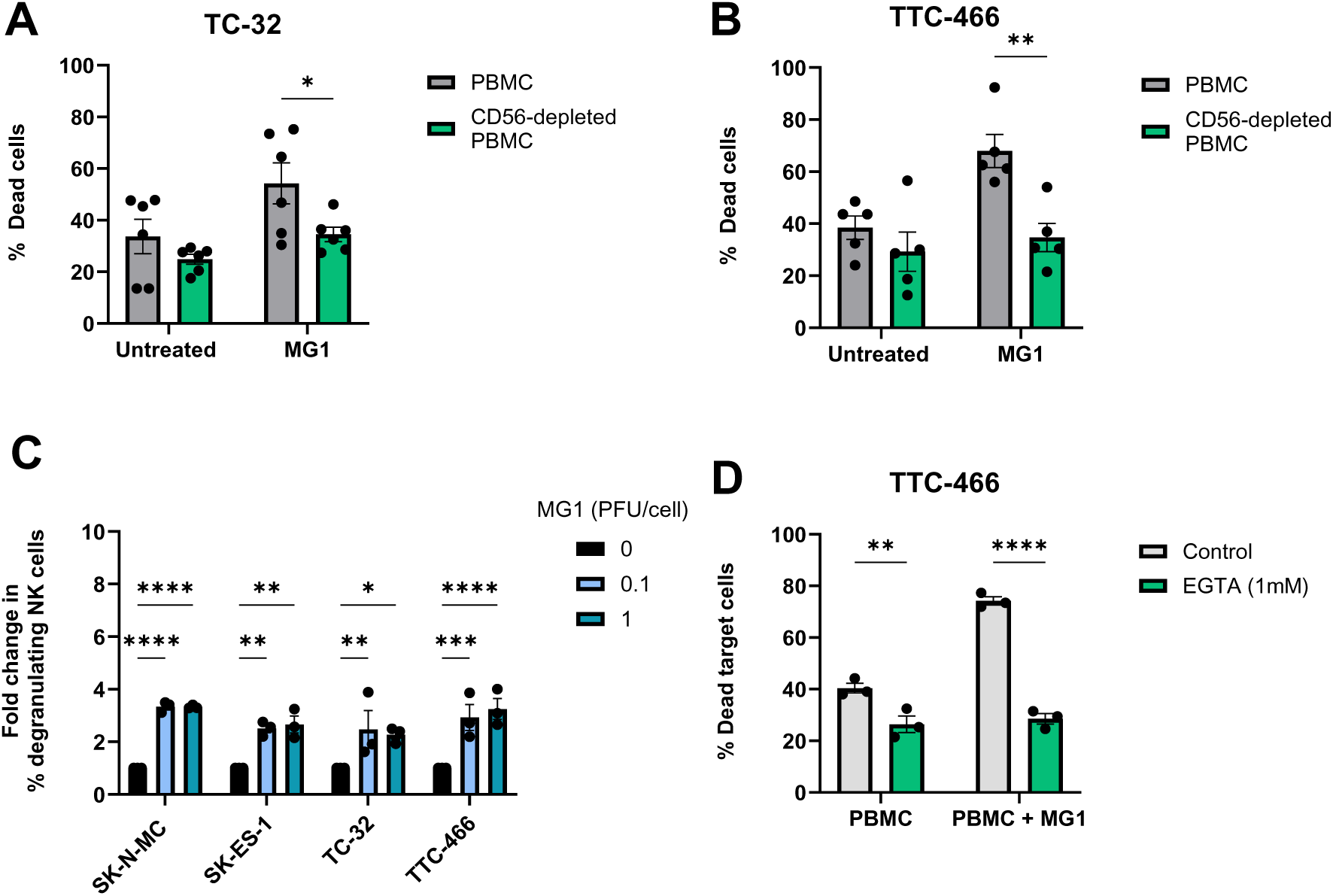
MG1 stimulates immune-mediated killing of EWS targets by NK degranulation. (A-B) NK cells were depleted from PBMC using CD56 magnetic bead selection and treated ± MG1 for 48 hours. Whole PBMC or CD56-depleted PBMC were co-cultured with cell tracker green stained (A) TC-32 and (B) TTC-466 EWS target cells at a ratio of 25:1 for 5 hours. Co-cultures were stained with LIVE/DEAD™ Fixable Yellow Dead Cell Stain and the percentage of dead target cells assessed by flow cytometry. (C) EWS cell lines spheroid cultures were generated over 7 days and PBMC were treated ± MG1 at 0.1 or 1 PFU/cell for 48 hours. PBMC were then co-cultured at a ratio of 10:1 with EWS spheroids for 4 hours. The percentage of degranulating NK cells was detected by staining with CD56, CD3 and CD107a antibodies and flow cytometry. Results presented as fold change in degranulating NK cells relative to untreated control. (D) PBMC were treated ± MG1 at 1 PFU/cell for 48 hours and then treated ± 1 mM EGTA for 30 minutes. PBMC were then co-cultured with TTC-466 EWS target cells at a ratio of 25:1 for 5 hours. Cells were stained with LIVE/DEAD™ Fixable Yellow Dead Cell Stain and the percentage dead target cells assessed using flow cytometry. All results presented are the mean ± SEM for a minimum of n=3 independent PBMC donors.

## Funding

The authors declare that financial support was received for the research, authorship, and/or publication of this article. T.B was funded by The Bone Cancer Research Trust PhD studentship (BCRT/6118) and National Institute for Health and Care Research (NIHR) Transatlantic Development and Skills Enhancement Award (NIHR304246). V.A.J was funded by Ovarian Cancer Action. E.A.R was funded by was supported by The Ewing’s Sarcoma Research Trust, The Bone Cancer Research Trust and FP7 grant “EURO EWING Consortium” (602856). R.T.B was funded by Sarcoma UK (SUK.10.2021). N.J.C. and J.K were supported by the Intramural Research Program of the National Cancer Institute (NCI), Center for Cancer Research (CCR), National Institutes of Health (NIH), Department of Health and Human Services (DHHS), USA: ZIA BC 011704 supports N.J.C; ZIA BC 010806 supports J.K. Other authors were supported by their respective institutions. The views expressed are those of the author(s) and not necessarily those of the NHS, the NIHR or the Department of Health and Social Care, or any other funders of this work.

## Acknowledgments

The authors would like to extend their gratitude to The Bone Cancer Research Trust for generous funding and support, which made this research possible. The commitment of The Bone Cancer Research Trust to advance the field of primary bone cancer research has been instrumental in allowing us to carry out this work. We also thank the other organisations who supported the study. We are indebted to Dr Lindy Visser and colleagues from the Princess Máxima Center for Pediatric Oncology, Utrecht, Netherlands for assistance with analysis of their Ewing sarcoma scRNAseq dataset.

## Author contributions

**TB:** Data curation, Formal analysis, Funding acquisition, Investigation, Methodology, Project administration, Resources, Supervision, Validation, Visualisation, Writing – original draft, review & editing. **VAJ:** Investigation, Methodology, Resources, Supervision, Writing – review & editing. **EAR:** Resources, Writing – review & editing. **RTB:** Investigation, Writing – review & editing. **MY:** Investigation, Writing - review. **HO:** Resources, Writing – review & editing. **DM:** Resources, Writing – review & editing. **PVG:** Resources, Writing – review & editing. **NJC:** Supervision, Resources, Writing – review & editing. **JK:** Resources, Writing – review & editing. **JCB:** Resources, Writing – review & editing. **SAB:** Resources, Writing – review & editing. **FEM:** Conceptualisation, Funding acquisition, Supervision, Writing – original draft, review & editing. **GPC:** Conceptualisation, Funding acquisition, Supervision, Writing – original draft, review & editing.

## Conflict of interest

The authors declare that the research was conducted in the absence of any commercial or financial relationships that could be construed as a potential conflict of interest.

## Notes

### Competing Interest Statement

The authors have declared no competing interest.

### Summary of Updates

The author list has been updated on this manuscript.

