## Supplementary Tables for "Oncolytic Maraba virus MG1 mediates direct and natural killer cell-dependent lysis of Ewing sarcoma"

**Supplementary Table 1: Cell lines and primary cell cultures**

| **Cell type** | **Origin** | **Details** | **Growth medium** |
| --- | --- | --- | --- |
| **Cell lines** | | | |
| SK-N-MC | Human Ewing sarcoma | EWSR1::FLI1 (type 1) fusion; Female; age 14 years | DMEM:F12 + 10% FBS |
| SK-ES-1 | Human Ewing sarcoma | EWSR1::FLI1 (type 2) fusion; Male; age 18 years | McCoys 5A + 10% FBS |
| TC-32 | Human Ewing sarcoma | EWSR1::FLI1 (type 1) fusion; Female; age 17 years | RPMI + 10% FBS |
| TTC-466 | Human Ewing sarcoma | EWSR1::ERG fusion; Female; 4 years | RPMI + 10% FBS |
| Vero | Kidney epithelial cells (African green monkey) |  | DMEM + 10% FBS |
| **Primary cell cultures** | | | |
| MSC | Human Bone marrow-derived mesenchymal stem cells | Male; age 22 years | StemMACS™ MSC Expansion Media +100 units of penicillin and 0.1 mg/mL streptomycin |
| CCRG1-L-017 | Human Ewing sarcoma | EWSR1::FLI1 (type 1) fusion; Male; age 15 years | RPMI + 10% FBS + 100 units of penicillin and 0.1 mg/mL streptomycin |
| CCRG1-L-023 | Human Ewing sarcoma | EWSR1::FLI1 (type 2); Male; age 11 years | RPMI + 10% FBS + 100 units of penicillin and 0.1 mg/mL streptomycin |
| CCRG1-L-066 | Human Ewing sarcoma | EWSR1::ERG; Female; age 17 years | RPMI + 10% FBS + 100 units of penicillin and 0.1 mg/mL streptomycin |
| Peripheral blood mononuclear cells | Human Healthy donor | Adult healthy donor blood from apheresis cones | RPMI + 10% FBS |

RPMI; Roswell Park Memorial Institute, DMEM; Dulbecco’s Modified Eagles Medium

**Supplementary Table 2**: **Antibodies, fluorescent stains, other reagents and buffers**

| **Flow cytometry antibodies** | | | | | |
| --- | --- | --- | --- | --- | --- |
| **Target protein** | **Target species** | **Isotype** | | **Fluorophore** | **Manufacturer** |
| CD3 | Human | Mouse IgG2aκ | | PerCP | Miltenyi Biotec |
| CD56 | Human | Mouse IgG1k | | eFluor450 | Thermofisher |
| CD107a | Human | Mouse IgG1k | | PE | Biolegend |
| CD99 | Human | Mouse  IgG2ak | | PE | Biolegend |
| LDLR | Human | Mouse IgG1 | | PE | R&D systems |
| CD69 | Human | Mouse IgG1k | | PE | Biolegend |
| CD317 | Human | Recombinant human IgG1 | | PE | Miltenyi Biotec |
| **ELISA antibodies** | | | | | |
| **Product** | | | **Manufacturer** | | |
| Anti-human IFN-α mAb (MT1/3/5), unconjugated | | | Mabtech | | |
| Anti-human IFN-α mAb (MT2/4/6), biotin | | | Mabtech | | |
| Human IFN-beta ELISA Kit - Quantikine | | | R&D systems | | |
| **Fluorescent stains** | | | | | |
| **Product** | | | **Manufacturer** | | |
| CellTracker^TM^ Green CMFDA Dye | | | Thermofisher | | |
| LIVE/DEAD® Fixable Yellow Dead Cell Stain  Kit | | | Thermofisher | | |
| **Other reagents** | | | | | |
| **Product** | | | **Manufacturer** | | |
| CellTiter-Glo® reagent | | | Promega | | |
| Brefeldin A | | | Biolegend | | |
| Lymphoprep | | | StemCell Technologies | | |
| Foetal Bovine Serum | | | Sigma | | |
| Methylthialazole Tetrazolium; MTT | | | Sigma | | |
| CD56 Microbeads Human | | | Miltenyi Biotec | | |
| Accutase | | | Thermofisher | | |
| EDTA | | | Sigma | | |
| Human IFNβ recombinant protein | | | PeproTech | | |
| **Buffers** | | | | | |
| MACS buffer | | | PBS + 1% FBS + 0.4% 0.5M EDTA | | |
| FACS buffer | | | PBS + 10% FBS + 0.1% sodium azide | | |
